# Learning induces unique transcriptional landscapes in the auditory cortex

**DOI:** 10.1101/2023.04.15.536914

**Authors:** G. Graham, M.S. Chimenti, K.L. Knudtson, D.N. Grenard, L. Co, M. Sumner, T. Tchou, K.M. Bieszczad

**Affiliations:** Neuroscience Graduate Program, Rutgers Univ., Piscataway, NJ; Behavioral and Systems Neuroscience, Dept. of Psychology, Rutgers Univ., Piscataway, NJ; Iowa Institute of Human Genetics, Univ. of Iowa Carver College of Medicine, Iowa City, IA; Rutgers Center for Cognitive Science, Rutgers Univ., Piscataway, NJ; Dept. of Otolaryngology-Head and Neck Surgery, Rutgers Robert Wood Johnson Medical School, New Brunswick, NJ

## Abstract

Learning can induce neurophysiological plasticity in the auditory cortex at multiple timescales. Lasting changes to auditory cortical function that persist over days, weeks, or even a lifetime, require learning to induce *de novo* gene expression. Indeed, transcription is the molecular determinant for long-term memories to form with a lasting impact on sound-related behavior. However, auditory cortical genes that support auditory learning, memory, and acquired sound-specific behavior are largely unknown. This report is the first to identify in young adult male rats (Sprague-Dawley) genome-wide changes in learning-induced gene expression within the auditory cortex that may underlie the formation of long-lasting discriminative memory for acoustic frequency cues. Auditory cortical samples were collected from animals in the initial learning phase of a two-tone discrimination sound-reward task known to induce sound-specific neurophysiological and behavioral effects (e.g., Shang et al., 2019). Bioinformatic analyses on gene enrichment profiles from bulk RNA sequencing identified *cholinergic synapse (KEGG 04725), extra-cellular matrix receptor interaction (KEGG 04512)*, and *neuroactive ligand-receptor interaction (KEGG 04080)* as top biological pathways for auditory discrimination learning. The findings characterize key candidate effectors underlying changes in cortical function that support the initial formation of long-term discriminative auditory memory in the adult brain. The molecules and mechanisms identified are potential therapeutic targets to facilitate lasting changes to sound-specific auditory function in adulthood and prime for future gene-targeted investigations.

---

A well-accepted concept in the field of learning and memory is that memories are stored where they are processed (Nadel & Hardt, 2011). Biological events known as memory consolidation can stabilize transient neural representations evoked by a sensory experience (Lechner, Squire, & Byrne, 1999; McGaugh, 2000; Dudai, 2012). Fundamental and evolutionarily conserved mechanisms that initiate memory consolidation are transcription and translation, defined as the active expression of genes and their subsequent protein products, respectively (Alberini & Kandel, 2014; Costa-Mattioli et al., 2009). We hypothesized that sound discrimination learning induces *de novo* gene expression within the auditory cortex. Highly specific representations of sound cues can outlast the transience of experience (seconds and minutes) by consolidating into long-term memory (hours to days) that later guides sound-cued behavior. While distinctive regional transcriptomic profiles are thought to be responsible for memory consolidation (Katzman, et al., 2021), learning-induced transcription events that support memory formation in the central auditory system are severely under-described. In contrast, the auditory cortex has been exceptionally well-described in studies of auditory learning and memory at the level of *neurophysiological* changes particularly in receptive fields and tonotopic maps (Schreiner & Polley, 2014; Weinberger N. M., 2015; Pienkowski & Eggermont, 2011), cortico-cortical and cortico-fugal connectivity (Souffi et al., 2021; Lesicko & Geffen, 2022; Schreiner & Polley, 2014; Liu et al., 2011; Xiong, Znamenskiy, & Zador, 2015), including at multiple time scales (Froemke & Martins, 2011; Froemke & Schreiner, 2015; Fritz, Elhilali, David, & Shamma, 2007; Tchernichovski & Margoliash, 2013). Moreover, neurophysiological changes are linked to behavior, e.g., for cue-directed action (Letzkus, Wolff, & Lüthi, 2015), attention (Fritz et al., 2007; Elhilali et al., 2007) and memory for sound signals (Bieszczad & Weinberger 2010; Grosso et al., 2015; Aschauer & Rumpel, 2016; Concina, Renna, Grosso, & Sacchetti, 2019; Letzkus, Wolff, & Lüthi, 2015; Ghosh & Zador, 2021). Decades of evidence for learning-induced neurophysiological plasticity in the auditory cortex has shown shared behavioral characteristics of auditory memory (Weinberger 2007a; 2007b), making it a top candidate region for auditory memory consolidation. By investigating auditory cortex bioinformatically, we also leverage an unbiased opportunity to uncover brain-wide common or distinct biological gatekeepers of neuroplasticity underlying adaptive auditory function. From a broader perspective, regional transcriptomic profiling can lead to a more complete understanding of how the sensory system is modified by experience in the service of memory. Learning-induced transcriptomic profiles identified within the auditory cortex can validate, extend, or lead to new biological models of processes that support or impair adaptive auditory processing. The consolidation of auditory memory is likely a product of experience-dependent changes in gene expression that affect cellular functions within the auditory cortex, which in turn promote lasting changes to sound-evoked neurophysiological responsivity that alter sound-cued behaviors.

To identify learning-induced transcripts, expression profiling analysis, using RNA sequencing (RNAseq) was performed on samples of anatomically defined auditory cortex (Bregma -3.10 mm, Interaural 6.90 mm; Paxinos & Watson, 2007) from water-restricted adult rats trained to bar press to pure tones for water rewards. Responses to a target pure tone frequency (5.0 kHz; 65 dB SPL) resulted in reward, while responses to the non-target (11.5 kHz; 65 dB SPL) were unrewarded and initiated a time-out period that extended the time to the next trial. This two-tone discrimination task (2TD) is not difficult perceptually; the acoustic frequencies are over an octave apart and easily distinguished by rodents (Talwar & Gerstein, 1998). Rather, the behavioral challenge was associative in nature: 2TD performance demands memory for which tone is associated with reward (vs. no reward). Trained rats (N=8) were sacrificed after three consecutive daily 45-minute 2TD training sessions and compared to a group of sound-naïve rats (N=4). This is a time point early in training when animals are still acquiring the 2TD task. Early task acquisition was targeted to capture initial learning-induced transcriptional events that set the stage for later increases in 2TD performance observed in behavior over weeks of training. For example, the average performance was above chance, but only 66±7.81% after 3 days, compared to ≥90% after extended training (Shang, Bylipudi, & Bieszczad, 2019). To capitalize on an opportunity to identify the expression of genes relevant for successful sound-specific associative memory, we leveraged an HDAC-inhibitor that targets epigenetic regulation of activity-dependent gene expression in learning and memory formation (McQuown, et al., 2011; Kwapis, et al., 2017; Malvaez, et al., 2012). Importantly, a decade of work has shown that HDAC-inhibition can facilitate learning-induced neurophysiological plasticity in sound-evoked auditory cortical responses (relative to vehicle) and enhances auditory discriminative behaviors (Bieszczad et al., 2015; Phan et al., 2017; Shang, Bylipudi, & Bieszczad, 2019; Rotondo & Bieszczad, 2020; Rotondo & Bieszczad 2021a; 2021b) including in humans (Gervain, et al., 2013). Half of the trained animals were treated with the HDAC-inhibitor, RGFP966 (N=4; 10 mg/kg, ApexBio, cat#A8803; *sub*.*cu*. injection), while the other half were identically trained but administered vehicle solution (N=4; matched volume, *sub*.*cu*. injection; **Fig. 1a**). There were no differences in 2TD performance between groups at the time of brain collection (RGFP966: 61±5.0% vs. Vehicle: 71±7.0%; *t*_(5.9166)_ = -1.21, p=0.272; Welch’s t-test). Brains were promptly collected and flash-frozen at a timepoint consistent with the peak concentration of the inhibitor in auditory cortex, one hour after the post-session injection of either the HDAC-inhibitor or vehicle (Bieszczad et al., 2015).

**Figure 1.**
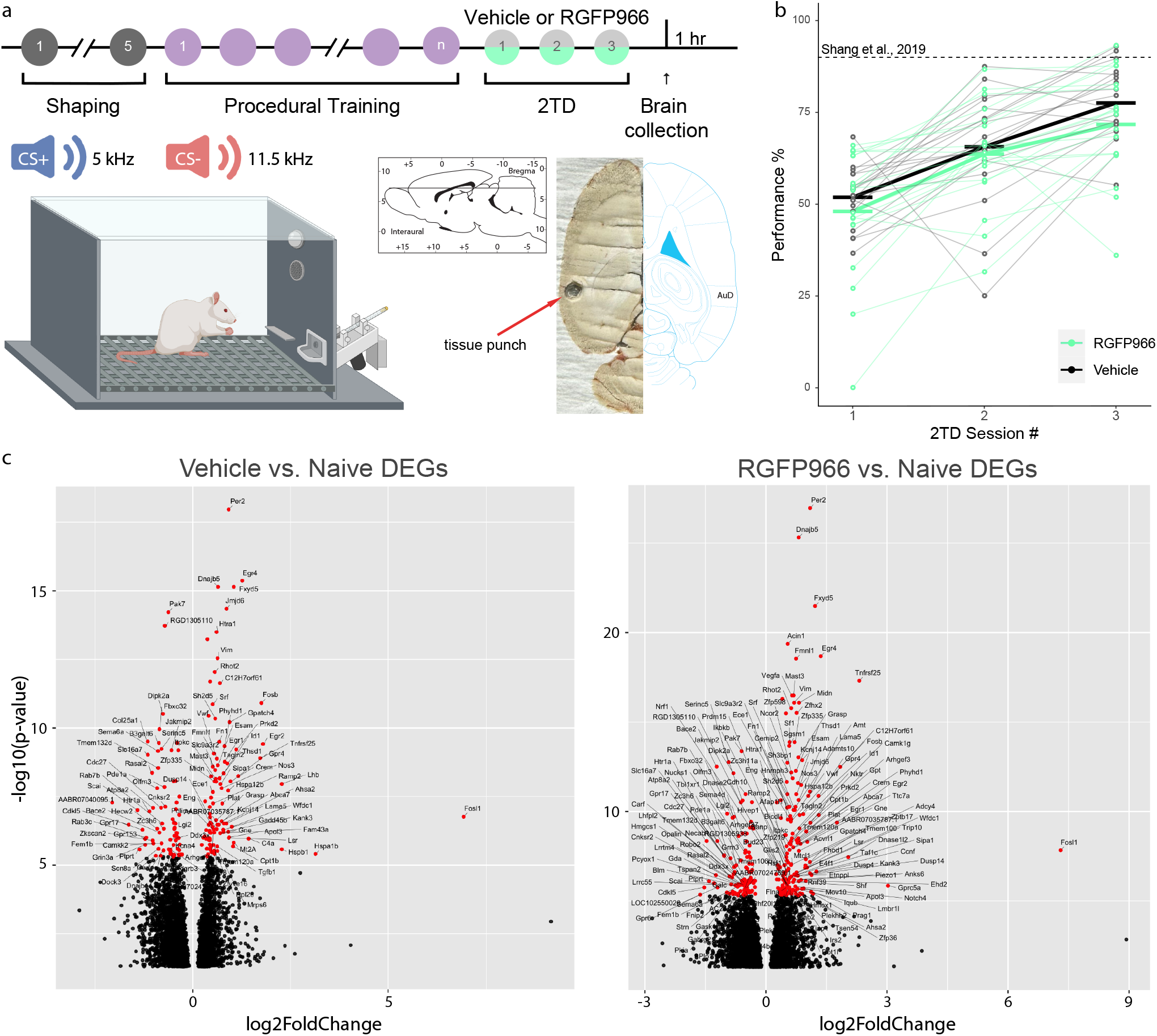
Behavioral timeline for tissue collection and use in RNA-sequencing. **(a) Shaping:** Rats were water restricted and trained to bar press for a water reward in single daily sessions over 5 days. **Procedural Training:** To put bar-pressing behavior under the control of sound, all rats were trained in daily 45-minute sessions to bar-press to broad-band noise for water rewards to a performance criterion of 80% correct, which all subjects achieved over 1-2 weeks with individual differences in the rate of acquisition. Thus, pairs of rats that similarly acquired the procedural aspects of the task were assigned to different treatment groups in the next phase of training (with the selective HDAC inhibitor, RGFP966, vs. Vehicle) so that both groups were matched in prior auditory experience and basic learning ability (as in Shang et al., 2019). **Two-tone discrimination (2TD):** Rats were trained in single daily 45-minute 2TD sessions, each immediately followed by injections of the HDAC-inhibitor, RGFP966, or Vehicle for 3 consecutive days. 2TD required rats to bar-press to a rewarded cue (5.0 kHz; 65 dB SPL) for water reward, and to avoid responding to the unrewarded cue (11.5 kHz; 65 dB SPL) (see inset). **Brain collection:** One hour after the last injection of RGFP966/Vehicle on the third day of 2TD training, rats were sacrificed for brain collection. Whole brains were rapidly removed and immediately flash frozen for storage at -80° C. To prepare for cryosectioning, brains were encased in optimal cutting temperature compound (OCT) and then stored at -20° C for 12-24 hours. Brains were sliced horizontally in a cryostat (Leica CM 3050S) at a thickness of 250 μm in four adjacent slices targeting auditory cortex (see inset) to collect 2 mm^3^ of cortical tissue from each hemisphere using a 1 mm round tissue micropunch (Harris). Thus, a total of 4 mm^3^ per brain region per subject was used to produce a single sample for subsequent RNA extraction and sequencing. **(b)** Plot showing performance values for each day of two tone discrimination (2TD). **(c)** Volcano plot showing auditory cortical learning-induced gene levels. Learning-induced genes were identified in transcript level comparisons between 2TD-trained vehicle-treated rats and sound-Naïve rats. Each dot represents a gene; some are labeled with gene names or identifiers. Significantly expressed genes (red) were determined by a set threshold of -log10(p-value) of 5.301 (*y-axis*; actual adjusted p-value = 0.000005). Genes included in enrichment analyses were only those that were significantly expressed and passed a log2FoldChange threshold of 0.65 (*x-axis*; actual fold-change value greater than 1.5692).

The findings herein are the first to identify a transcriptomic profile of associative learning within the auditory cortex. Learning in the two-tone discrimination task induced changes in the transcription levels of hundreds of genes (compared to sound-naïve) (**Fig. 1b**). A hierarchical clustering algorithm showed that genes are uniquely upregulated or downregulated, revealing a complex network of cortical gene expression events induced by auditory learning (**Fig. 2a**). Enrichment analyses (iPathwayGuide™; Impact Analysis method) identified the *cholinergic synapse* (differentially expressed genes (DEG) / all genes (ALL): 22/101; p = 0.004, Bonferroni correction) as the top biological pathway (**Table 1**). This result is consistent with research since the 1990s highlighting the sufficiency of cholinergic signaling in the auditory cortex for neurophysiological plasticity and related auditory behavior (Froemke & Martins, 2011; Weinberger, 2003; Bakin & Weinberger, 1996; Kilgard & Merzenich, 1998). Transcript levels induced by learning under HDAC-inhibition were either amplified in the same direction (by further increases or further decreases in unique gene transcript levels) (**Fig. 2b**), or blunted compared to learning alone (**Fig. 2c**). One top biological pathway under these conditions was the *Neuroactive ligand-receptor interaction* (DEG/ALL: 30/194; p = 0.016; **Table 1**) which involved effectors crucial to neural activation. Another top pathway was the *extracellular matrix-receptor (ECM-receptor) interaction* (DEG/ALL: 14/69; p = 0.04; **Table 1**). Components of the ECM are critical for cortical plasticity and memory consolidation (Happel, et al., 2014; Banerjee, et al., 2017; El-Tabbal, et al., 2021; Sonntag, et al., 2015). These pathways offer potential alternatives, perhaps complementary, to cholinergic signaling that can facilitate lasting experience-dependent changes in auditory function (Ji, Gao, & Suga, 2001; Luo & Yan, 2013; Metherate, 2011). Other genes whose expression changed with auditory learning but no further with HDAC-inhibition were likely related to procedural conditions of the task, rather than to sound-specific associative memory. For example, the *apelin signaling pathway* was identified (DEG/ALL: 20/116; p = 0.0005), which is important in the brain for the homeostatic regulation of water intake (Hu, et al., 2021). A direct comparison of transcript levels between the two groups of trained rats (learning with or without HDAC-inhibition) showed very few uniquely differentially expressed genes (DEGs). A threshold appreciably used to identify the most likely DEGs (p = 0.05) found only *Adamts13, U6, Rexo4*, and *Cabin1* were differentially expressed (**Fig. 2d**). Thus, the major effect of HDAC-inhibition appears to modulate the expression of genes induced by auditory learning under normal conditions, rather than to newly recruit unique genes to subserve sound-specific memory. Together, these findings show large-scale transcriptomic changes occur in the auditory cortex early in training as adult animals learn to discriminate associative relationships between sound cues. Together with established neurophysiological and behavioral reports of HDAC-inhibition to promote auditory function, these findings support the tactic of using an HDAC-inhibitor to distinguish genes whose expression determines the success of auditory memory formation.

**Table 1.**
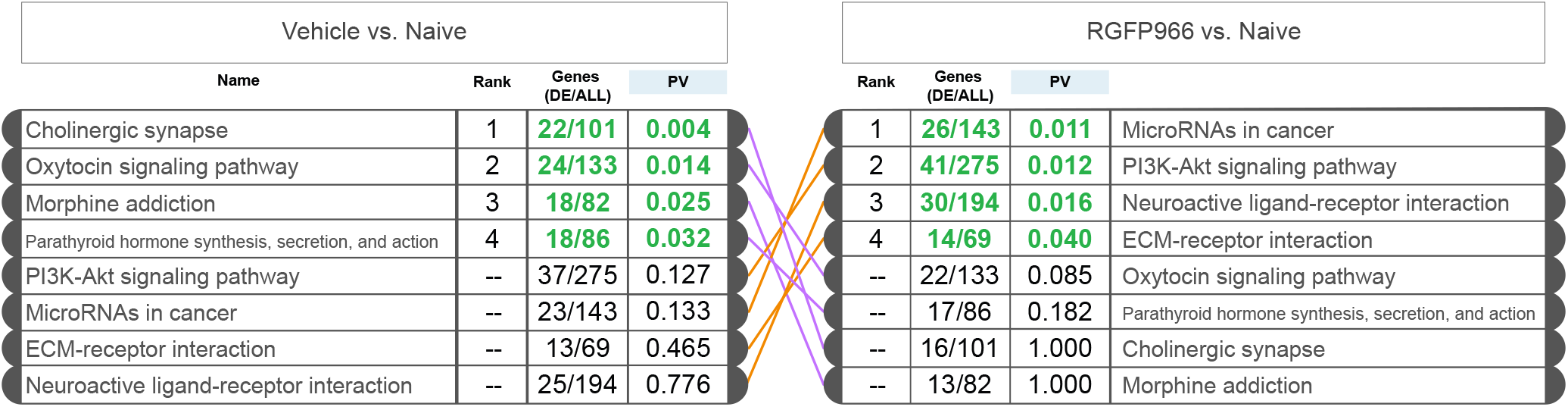
Enrichment analysis of learning-dependent transcriptional changes identifies potential downstream pathways important for auditory function. A meta-analysis shows the rank list order of unique biological pathways involved in learning-induced transcriptional changes identified in iPathwayGuide© (“Learning-induced”: Vehicle vs. Naïve; “Learning-induced with RGFP966”: RGFP966 vs. Naïve). Ranks are assigned by p-value (with Bonferroni correction) to identify top pathways. The number of genes reported as DEGs (DE) out of all genes involved in a pathway according to the iPathwayGuide© databases (ALL) are listed as “DE/ALL”. *PV: p-value*.

**Figure 2.**
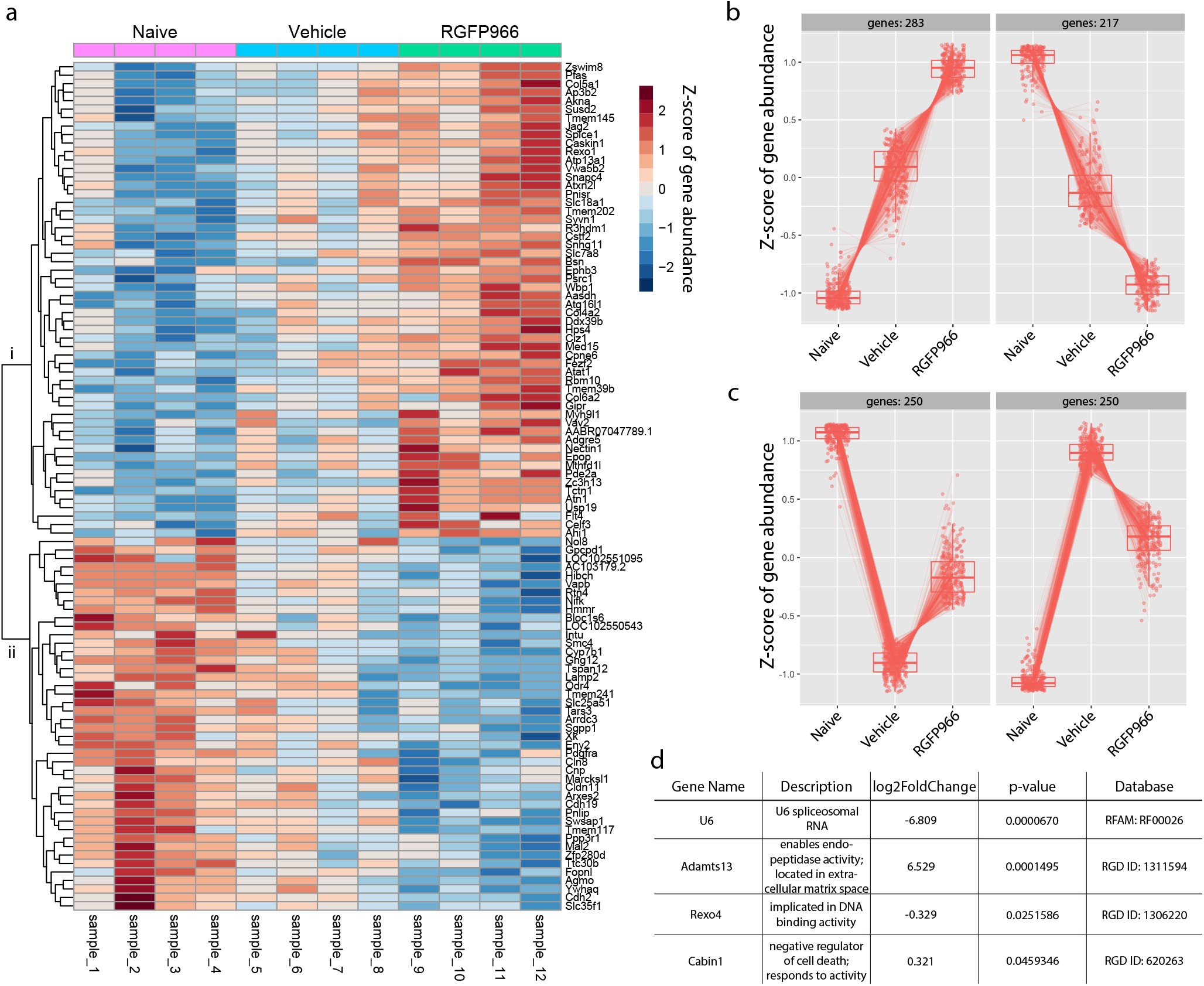
RNA-sequencing reveals abundant transcriptional changes in auditory cortex. RNA-sequencing of individual subject samples of auditory cortical tissue from naïve (N=4), or 2TD-trained vehicle treated (“Vehicle”: N=4), and 2TD-trained HDAC-inhibitor treated (“RGFP966”: N=4) rats. **(a)** Heatmap of 100 genes with the most different levels of expression between trained rats treated with RGFP966 vs. Naïve. Transcript abundance levels for these 100 genes were z-scored with Naïve transcript levels set to -1, and genes with largest z-score relative to -1 were analyzed and shown for all three groups. A hierarchical clustering algorithm applied to categorize relative transcript levels across the 3 groups revealed two major clusters of effects: genes that are *upregulated* with 2TD learning and further upregulated by adding HDAC-inhibition (RGFP966) to 2TD learning (denoted as “i” in cluster bracket on left of the heatmap plot), as opposed to genes that are *downregulated* with learning and further down by adding HDAC-inhibition to 2TD learning (denoted as “ii”). **(b)** Box and whisker plots show the same analysis extended to 500 genes with the largest difference in z-scores between RGFP966 and Naïve groups. Each dot represents relative gene abundance of the same gene across the 3 conditions (Naïve, Vehicle or RGFP966). The left panel shows that 283/500 of the genes identified are upregulated with 2TD learning and further upregulated by adding HDAC-inhibition, while the right panel shows 217/500 genes are downregulated with learning and further downregulated by adding HDAC-inhibition. **(c)** Box and whisker plots of the top 500 genes with the largest difference in z-scores between Vehicle and Naïve groups. As in (b), each dot represents relative gene abundance of the same gene across the 3 conditions (Naïve, Vehicle or RGFP966). The left panel shows that 250/500 of genes identified are *downregulated* with 2TD learning (Vehicle), which is blunted by adding HDAC-inhibition (RGFP966). The right panel shows that the other 250/500 genes identified are *upregulated* with 2TD learning (Vehicle), which is blunted by adding HDAC-inhibition (RGFP966). **(d)** A direct comparison of differentially expressed genes (DEGs) between 2TD-trained groups, Vehicle vs. RGFP966, identified only four DEGs determined with an adjusted threshold of p<0.05.

Several genes of interest (GOIs) were selected from the enrichment analyses in biological pathways to validate genome-wide sequencing with a gene-targeted approach. Samples were collected from separate cohorts that replicated the two groups of trained and treated animals at the same early-in-training timepoint (i.e., one hour after the third 2TD training session) for use in gene expression analysis using quantitative real-time polymerase chain reaction (qRT-PCR). There were no 2TD performance differences in replicate cohorts (RGFP966_1_: 61±5.0% vs. RGFP966_2_: 67±6.0%; *t*_(8.441)_ = -0.834, p = 0.4274; and Vehicle_1_: 71±6.0% vs. Vehicle_2_: 74±3.0%; *t*_(4.0809)_ = -0.444, p = 0.6796; Welch’s t-test). The first GOI was *Egr1*, a well-studied, activity-dependent “immediate-early gene” known to peak within the first hours of learning and critical for memory formation (Duclot & Kabbaj, 2017). It was a top learning-induced gene with confirmed increased expression that was likewise amplified with HDAC-inhibition in the separate cohort of animals in gene-targeted study (**Fig. 3**). The same was true for *Per2*, a gene involved in the regulation of circadian rhythms and serotoninergic pathways in the brain (Bae, et al., 2001; Albrecht, et al., 2001; Cuesta, et al., 2009; Reh, et al., 2020). *Per2* is part of the “clock gene” family that has been linked to HDAC modulation in other brain regions (Kwapis, et al., 2018) and may have a role in auditory function as well (Reh et al., 2020). In contrast, some GOIs identified genome-wide were only partially confirmed. *Chrna7* encodes a subunit of the dynamic nicotinic acetylcholine receptor (nAChR) that was found to be downregulated with auditory learning, consistent with neurophysiological evidence (Takesian, et al., 2018; Kuchibhotla, et al., 2017). Thus, we expected the gene-targeted study to confirm *Chrna7* down-regulation in all 2TD trained animals, but down-regulation was apparent only in vehicle-treated animals that learned 2TD without HDAC-inhibition. Additional selected GOIs were from the *Nr4a* orphan nuclear receptor family, *Nr4a1* and *Nr4a2*, which are immediate early genes reported to be necessary for propagating the downstream effects of the selective HDAC target of RGFP966 in other brain regions (McQuown et al., 2011; Kwapis, et al., 2019). Consistent with prior reports of task-dependent regional differences in *Nr4a* family gene expression (McNulty, et al., 2012), *Nr4a1* but not *Nr4a2* was up-regulated in auditory cortex after auditory learning. Other DEGs, *Htr1a* or *Adamts13*, were not confirmed by gene-targeted study likely due to differences in known transcript variants not detected by our custom-designed gene-targeted probes, or due to type I error, or to subtle behavioral differences between trained cohorts of animals that went undetected in 2TD performance measures. We also investigated *Lynx1*, a gene with a long history of function for re-opening “critical period”-like plasticity in sensory cortex (Morishita et al., 2010), likely by its action on cholinergic and serotonergic modulation (Takesian, et al., 2018). However consistent with prior genome-wide reports that failed its detection (Kalish, et al., 2020), it was neither detected in our RNAseq dataset, nor in gene-targeted study. Though it remains a challenge to determine whether discrepancies can be explained by real biological or by individual variability between animals, or due to technical variability with low abundance or cell-type specific expressing transcripts, we report exceptionally consistent results for some GOIs. As the learning-induced expression of *Chrna7, Egr1*, and *Per2* is consistent between the gene-targeted and genome-wide approaches, and between different cohorts of trained animals, it stands to reason that these genes may be of the most fundamental players for lasting changes to auditory function and memory and are now prime for future investigation.

**Figure 3.**
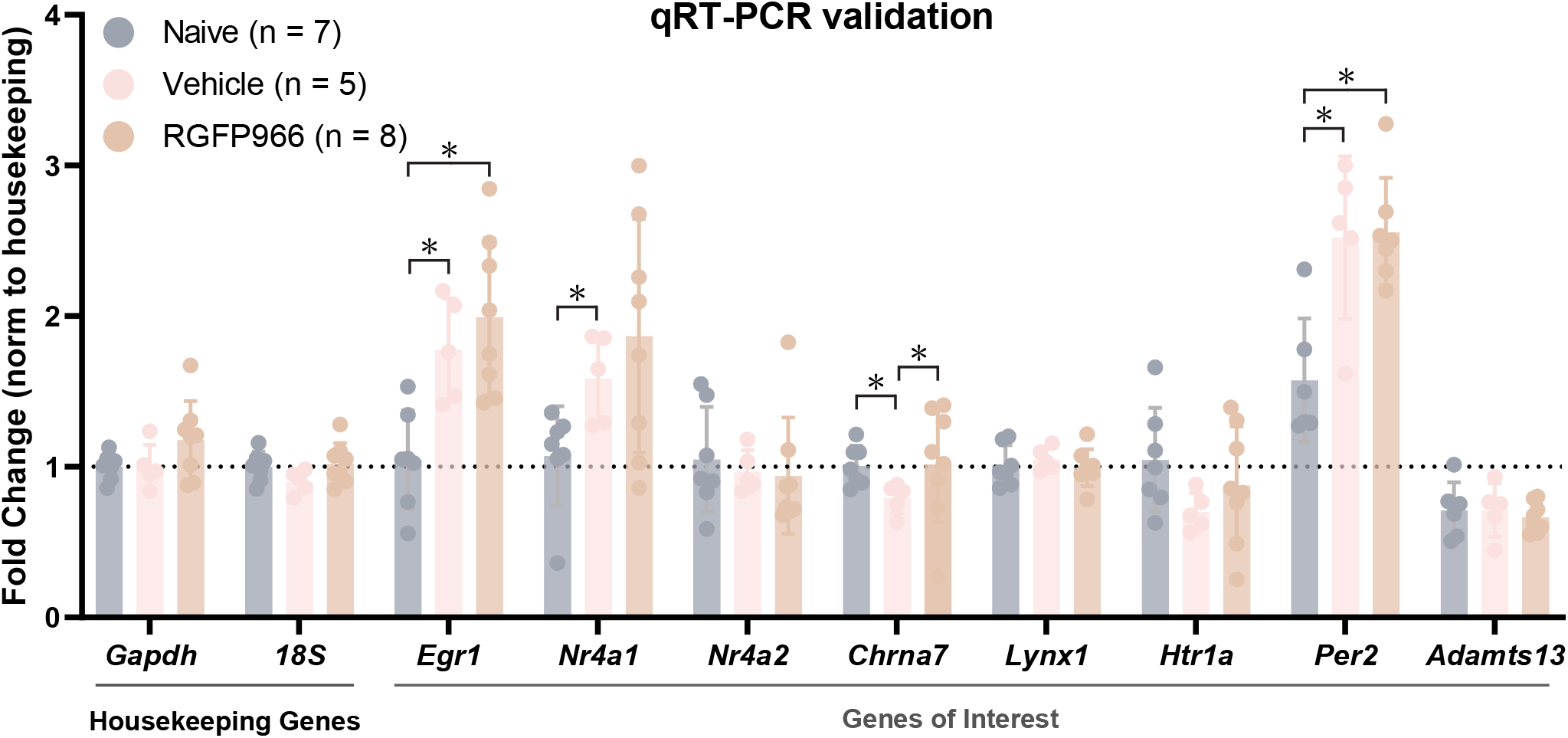
A gene-targeted approach using qRT-PCR validates selected genes of interest identified from genome-wide RNAseq. A second cohort of animals was used for qRT-PCR analysis. They were trained identically to the animals that generated samples for RNAseq (Naïve: N=7; Vehicle: N=5; RGFP966: N=8). As for RNAseq, brains were collected 1 hour post injection of RGFP966 after the third daily session of 2TD training to generate auditory cortical RNA samples. All values were normalized to the housekeeping genes *Gapdh* or *18S* (values plotted show normalization to *Gapdh*). qRT-PCR results demonstrate robust learning-induced increases in expression of *Egr1, Nr4a1*, and *Per2*. All data are expressed as means (±standard errors). *p<0.05 (Kolmogorov-Smirnov test).

Considered together, the findings reveal a dynamic transcriptional landscape in the adult auditory cortex that could support emergent auditory function in neurophysiology and behavior. Given the growing evidence of epigenetic controls on sensory system function (c.f., Shang & Bieszczad, 2022), it is exciting to consider how epigenetic mechanisms play a role in balancing stability with the experience-dependent plasticity of auditory cortical circuits. For example, the current genome-wide dataset also identified a learning-induced reduction in the expression of a repressive histone deacetylase, *Hdac9*, that was absent with HDAC-inhibition. It is tempting to liken the effect of reduced HDAC9 expression to the effect of pharmacologically inhibiting HDACs to promote learning-induced transcription. Another important class of epigenetic regulators alter DNA rather than histones. Examples are *Tet1* and *Gadd45b* that both impact DNA methylation (Bayraktar & Kreutz, 2018) and were found to be down- and up-regulated with learning, respectively, under HDAC-inhibition. Further, a class of microRNAs appeared as the top biological pathway induced by learning with HDAC-inhibition (see **Table 1**), which have gained traction in the field as key epigenetic regulators of neural transcriptional control (Saab & Mansuy, 2014). Higher-order molecular interactions between epigenetic players and the protein products of expressed genes are also likely. For example, the identified up-regulated *Egr1* can recruit *Tet1* to remove repressive methylation marks to activate downstream genes (Sun, et al., 2019). Further molecular studies will be needed to parse out the significance of interactions between epigenetic regulators in the auditory system during learning. Moreover, this report presents an immediate opportunity to establish links between select auditory cortical genes and their downstream effectors. Bridging the gap from these genes and molecules to their influences on sound-evoked neurophysiological events is essential to understand how the adult auditory system adapts with experience to alter behavior. Indeed, the translation of *de novo* transcripts are required for long-term changes to behavioral function because they produce lasting changes to cellular function. For example, key target effectors of learning-induced transcriptional processes may alter the availability of channels or receptors (Metherate, Intskirveli, & Kawai, 2012; Brown & Kaczmarek, 2011; Henton, Zhao, & Tzounopoulos, 2023) within circuits that determine sound-evoked threshold, responsivity, and receptive field architecture in the auditory system. It is reasonable to assume that different auditory tasks would recruit unique gene networks for biological pathways relevant to the particular cellular functions that would support learning for the task-relevant sound feature or task structure.

Overall, this report serves to act as a starting point to make RNAseq datasets from the learning, adult, auditory cortex available (see **Data Repository**). An effort to narrow the gap in knowledge exacerbated by the severe lack of studies that focus on molecular genetic processes in adult auditory cortex is now welcome. We encourage study beyond the spatio-temporal limitations of bulk RNA-sequencing whose sensitivity is limited technically also by read depth (Li & Wang, 2021), especially since modern *Omics* technologies are quickly evolving and improving. While research in the auditory cortex has made some initial strides in molecular genetics that have identified essential molecular cascades (Schicknick, et al., 2008), permissive IEG profiles (Mello, Velho, & Pinaud, 2006; de Hoz, et al., 2018; Peter, et al., 2011) and even chromatin dynamics (Peter, et al., 2021), there is precedent in the auditory periphery where transcription dynamics are beautifully studied (Kwan, 2016; Barta, et al., 2018; Li, et al., 2020; Ebeid, et al., 2017). Existing molecular tools like for single cell RNA sequencing (scRNAseq) and small-molecule fluorescent in situ hybridization (smFISH) will be useful to provide insights into experience-dependent cell-to-cell variation and molecular interactions within auditory cortical cells and circuits. Unlike bulk RNAseq, these methods honor the profound cellular diversity and higher order organization in auditory cortex, such as its layer-specific microcircuitry and lemniscal topography. Clever approaches to tag and sequence transcripts from only recently active auditory cortical cells (Cho, Huang, & Gray, 2016) may increase sensitivity enough to robustly obtain the most functionally relevant sets of genes from the most functionally relevant cell types and populations. Similar approaches could be used also subcortically to capture and contrast activity-dependent regional transcriptomic profiles in the cortex that honors the incredible integration of cortical with subcortical function of the auditory system under different listening conditions or behavioral demands. Fully characterizing cellular and regional distinctions in genetic and molecular mechanisms underlying auditory learning in the adult brain is paramount for developing site-selective and molecule-targeted precision therapeutics that enable robust and persistent functional changes to support specific hearing and listening abilities across the lifespan.

## Acknowledgements

The authors would like to thank Dr. Andrea Shang, Sean Tsaur, and Sooraz Bylipudi for their assistance in behavioral procedures and analysis; Dr. Mimi L. Phan for her assistance with RNA extraction and sample collection protocols; Dr. Troy A. Roepke, Ali Yasrebi, and Christopher O’Brien for their assistance with RNA sample purification analysis; and Alisa Ray for technical assistance. We would like to thank all current and former CLEF Lab personnel for their assistance and support.

This work was supported by the National Institutes of Health, National Institute of Deafness and Communication Disorders [R01-DC014753 to K.M.B.]; the School of Arts and Sciences at Rutgers University; the Aresty Foundation at Rutgers University with small grant funding for undergraduate research; and the Project SUPER program at Rutgers University with small grant funding for undergraduate research.

## Data Repository

Raw sequencing files are publicly available here: rutgers.box.com/s/x1oi6d3vh7q46niumyyvh3qs5c1zbqbe

