## Supplementary Methods for "Learning induces unique transcriptional landscapes in the auditory cortex"

*Subjects*: A total of 48 adult male Sprague-Dawley rats (250 – 300 g on arrival; Charles River Laboratories, Wilmington, MA) were used in behavioral and molecular experiments. All animals were individually housed in a temperature-controlled (24 ˚C) colony room on a 12-hour light/dark cycle. Subjects had ad libitum access to food and water prior to behavioral training. All procedures were approved and conducted according to guidelines by the Institutional Animal Care and Use Committee (IACUC) at Rutgers, the State University of New Jersey (Protocol No.: 999900026 (K.M.B)).

*Behavioral apparatus and sound stimuli*: All behavioral sessions were conducted in two identical instrumental conditioning chambers (H10-112TC-NSF; Coulbourn Instruments, Holliston, MA) within a sound-attenuated box. Daily training sessions were counterbalanced to ensure equal exposure to both chambers. Each chamber (12” W x 10” D x 12” H; wire mesh floor was fitted with a response lever (H21-03R), house light (H11-01R), a speaker (H12-01R), and a water delivery system (H14-05R). During training phases, animals could depress the response lever (“barpress”), which triggered the presentation of a water cup (~0.02cc) in the reward port (1.25” W x 1.625” H). A hand switch (H21-01) was used during early session to shape the animal’s barpress response to trigger presentations of the water cup that allowed access to the water reward. Behavioral responses were recorded using Graphic State 4 software (Coulbourn Instruments, Holliston MA) for offline analysis.

All auditory stimuli were generated using Tucker-Davis Technologies (TDT, Alachua, FL) and RPvdsEx software, and presented via the operant chamber’s wall-mounted speaker. White noise (during procedural training; **Fig. 1a**) was presented for 7 or 9 s in duration (75 dB SPL). Pure tones (during two-tone discrimination training; **Fig. 1a**) were always presented for 8 s (70 dB SPL). Sound levels were calibrated daily using a digital sound meter (Larson Davis SoundTrack LxT1).

*Behavioral training and pharmacological inhibition of HDAC3*: After one day of acclimating to the vivarium, rats were handled daily for a minimum of 3 days. Prior to the start of behavioral training, rats were placed on a schedule of restricted water until they reached 85% of the non-restricted weight of age-matched control animals. Water-restricted rats were then shaped inside sound-attenuated chambers to barpress for water rewards (**Fig. 1a**). Barpress shaping and subsequent sound training were as previously described (Shang, Bylipudi, & Bieszczad, 2019). Briefly, all animals were shaped to barpress over 5 days, and then trained in a procedural task to learn to bar-press to sound, which in this phase was a broad-band Gaussian noise stimulus (1-12.5 kHz band-pass filtered white noise; 75 dB SPL), to obtain a water reward. All animals learned successfully learned to associate this sound with reward prior to continuing to the next phase of training. Procedural training showed individual variability of how quickly the animals could learn the task to high levels of performance. Nonetheless, all animals were able to achieve a performance level of ≥90% for two consecutive days (mean = 92.85%, s.e.m. = 0.01%) after an average of 12.67 days (s.d. = 2.85 days). Performance was calculated using $Performance=\frac{\#BPs to white noise}{Total \#BPs} \times100\%$ .

The next phase of training was the two-tone discrimination (2TD) task (**Fig. 1a**), in which rats were trained to discriminate between two spectrally distinct sound frequencies. Barpresses to the S+ tone (5.0 kHz; 70 dB SPL) would result in the presentation of a water reward, while barpresses to the S- tone (11.5 kHz; 70 dB SPL) resulted in an error signal (flashing house light) and a “time-out” (an additional 6 s wait for the start of the next trial). S+ and S- trials were randomized and lasted for 8 s. Inter-trial intervals (ITIs) were on average 15 s (range: 5-25 s, randomized). Barpresses during a silent ITI were inconsequential (no time-out, no error signal, nor water reward). Daily sessions were 45 minutes in length. All animals performed the 2TD task for a total of three consecutive days. Rats were paired by a trained observer so that animals with similar rates of acquiring the procedural task were assigned different treatment conditions in the 2TD phase. Performance-matched pairs were received systemic injections of either the pharmacological class I HDAC3 inhibitor RGFP966 (Abcam Inc., ab144819; 10 mg/kg; s.c.; N = 12) or vehicle solution (N = 9) immediately after each 2TD session. Performance of the 2TD task was calculated as previously described: $Performance=\frac{{BP}^{S+}}{{BP}^{S+}+ {BP}^{S-}} \times100\%$(Shang, Bylipudi, & Bieszczad, 2019).

*Tissue collection and isolation of RNA*: One hour after animals received their third and final injection of RGFP966 after the third session of 2TD, brains were quickly dissected and flash-frozen in a beaker of 2-methylbutane placed upon dry ice. A single time window was chosen for brain collection and RNA-sequencing so that all samples were obtained from animals who had received an equal amount of exposure to sound and to treatment with injections of either RGFP966 (the pharmacological HDAC-inhibitor) or Vehicle. Brain collection occurred during the hours of 12:00 PM – 7:00 PM across groups and for trained and untrained animals. Flash-frozen brains were then stored at -80˚ C until future processing. To prepare brains for cryosectioning, brains were encased in optimal cutting temperature compound (OCT) and stored at -20˚ C for 12-24 hours to ensure brain tissue would reach -20˚ C for cutting. Brains encased in OCT would then be sliced horizontally in a cryostat (Leica CM 3050S) at a thickness of 250 µm. Using a 1 mm round tissue micropunch, 2 mm^3^ of auditory cortical tissue from each hemisphere (combined into one sample for a total of 4 mm^3^ per brain region) was sampled for RNA extraction. Auditory cortex location was identified using Paxinos and Watson’s *The Rat Brain in Stereotaxic Coordinates* (6th edition) as a reference (approximately A/P = -3.6 to -5.8 mm, M/L = ±6.4 mm D/V = -4.2 to -5.8 mm; and using hippocampus as landmark).

Total RNA from each sample was isolated using the PureLink^TM^ RNA Mini Kit (ThermoFisher) using the manufacturer’s protocol. RNA samples were then purified with the RNA Clean and Concentrator^TM^ -25 Kit (Zymo Research).

*General methods for gene expression analysis:* RNA-seq analysis was conducted by the Iowa Institute of Human Genetics (IIHG; Iowa City, IA, USA). Briefly, using 500 ng total RNA (all RIN values >8), sequencing libraries were generated using the Illumina TruSeq® Stranded mRNA Library Prep kit according to the manufacturer’s recommended protocols. Libraries were pooled and sequencing was performed on an Illumina NovaSeq 6000 running 100 bp paired-end SBS chemistry. Reads were processed with the ‘bcbio-nextgen.py’ open-source informatics pipeline developed primarily at Harvard Chan Bioinformatics (v.1.2.4) running on the Argon HPC resource at the University of Iowa. This pipeline includes ‘best practices’ approaches for read quality control, read alignment and quantitation. The ‘bcbio-nextgen.py’ pipeline was run in “RNA-seq” mode with the ‘rn6’ key as the selected genome build (internally referencing Ensembl assembly and genebuild ‘Rnor_6.0’). The pipeline aligned reads to the Rnor_6.0 genome using the splice-aware, ultra-rapid hisat2 aligner (2.2.1) and concurrently quantified reads to the transcriptome using the ‘salmon’ (1.4.0) aligner. Qualimap (2.2.2), a computational tool that examines hisat2 BAM alignment files, was used to examine the read data for quality control. Sequence quality scores passed basic checks and sequence duplication rates were within acceptable parameters. All samples passed QC for read alignment to exonic regions. Salmon-derived transcript quantifications (TPM or “transcripts per million”) were imported and summarized to estimated counts at the gene level using tximport (1.12.3) in Rstudio, as described in the best-practices DESeq2 vignette (https://bioconductor.org/packages/release/bioc/vignettes/DESeq2/inst/doc/DESeq2.html). Genes with fewer than 5 estimated counts across all samples were pre-filtered from downstream analysis, as per recommended procedure.

*Differential gene expression analysis for Gene Ontology (GO) biological process analysis, and gene set enrichment analysis (GSEA)*: Differentially expressed gene (DEG) analysis was conducted with DESeq2 (v.1.24.0) on estimated gene-level counts. An FDR of 5% and X < abs(logFC) < 10 was set as a cutoff for differential expression (DEGs). Heatmaps, line graphs, and volcano plots were generated using clusterProfiler packages in R/Bioconductor. Gene level, pathway, and DEG analyses were generated using iPathwayGuide (Advaita Bioinformatics, https://www.advaitabio.com/ipathwayguide; last accessed November 15, 2022). iPathwayGuide scores pathways using the Impact Analysis method (Draghici et al., 2007; Tarca et al., 2009, Khatri et al., 2007). Impact analysis uses the over-representation of DEGs (Bonferroni-corrected for family-wise error rate) in a given pathway and the perturbation of that pathway computed by propagating the measured expression changes across the pathway topology. The underlying pathway topologies, comprised of DEGs and their directional interactions, are obtained from the Kyoto Encyclopedia of Genes and Genomes (KEGG) database (release 100.0+/11-12, Nov 21) using the protein-coding rat genome as a reference set (Kanehisa et al., 2000; Kanehisa et al., 2010; Kanehisa et al., 2012; Kanehisa et al., 2014).

*Gene level analysis*: Transcript level abundance was analyzed by Z-scoring the TPMs of each gene. Only significantly modulated genes are included in these analyses (only the top 500 of 992 DEGs identified out of 14,196 genes with measured expression in the RGFP966 vs. Naïve comparison and only the top 500 of 958 DEGs identified out of 14,196 genes with measured expression in the Vehicle vs. Naïve comparison). Z-scores from Naïve transcripts were compared to Z-scores from RGFP966 and Vehicle transcripts shown in Figure 2a-c. Heatmaps, line graphs, and volcano plots were generated using clusterProfiler packages in R/Bioconductor. Dendrograms and hierarchical clustering of gene expression for heatmap generation were achieved in R by using the pheatmap package and function with Euclidean distance as a similarity measure and a complete-linkage clustering method (cluster_rows = TRUE, clustering_distance_rows = “euclidean”, clustering_method = “complete”).

*RNA-seq data validation with qRT-PCR*: RNAseq data were validated by qRT-PCR on mRNA extracted from distinct cohorts of rats. Five transcripts were chosen from a list of novel DEGs found enriched in each region at baseline. Three of the genes were hypothesis-driven based on auditory cortical physiological and behavioral literature, while another three genes were novel genes identified from gene level and DEG analyses in a comparison between Drug and Vehicle groups. All primer sequences were selected either from previous literature on brain tissue in rats (sequences provided in table below) or designed using NCBI Primer Blast and validated for target specificity by assessing melt-curves and PCR amplicon product sequencing. qRT-PCR analyses were performed in a QuantStudio 3 (Applied Biosystems) with SsoAdvanced^TM^ Universal SYBR Green Supermix (Bio-Rad). For each sample, 500 ng of cDNA was amplified using an iScript^TM^ cDNA Synthesis Kit (Bio-Rad).

| Gene | Forward primer | Reverse primer | Reference |
| --- | --- | --- | --- |
| *Gapdh* | GTGGACCTCATGGCCTACAT | TGTGAGGGAGATGCTCAGTG | Malvaez et al., 2018 |
| *18S* | CGGACAGGATTGACAGATTG | CAAATCGCTCCACCAACTAA | Chang et al., 2016 |
| *Egr1* | AACCCTACGAGCACCTGAC | CGGGTAGTTTGGCTGGGATA | Malvaez et al., 2018 |
| *Nr4a1* | CCGGTGACGTGCAGCAATTTTATGAC | GGCTAGAATGTTGTCTATCCAGTC | Malvaez et al., 2018 |
| *Nr4a2* | ATTGCTGCCCTGGCTATGGT | GACCATCCCATTATTGAAAGTCACATGGTC | Malvaez et al., 2018 |
| *Chrna7* | TGCACGTGTCCCTGCAAGGC | GTACACGGTGAGCGGCTGCG | Bychkov et al., 2018 |
| *Lynx1* | ACCACTCGAACTTACTTCACC | ATCGTACACGGTCTCAAAGC | Bychkov et al., 2018 |
| *Htr1a* | GACCACGGCTACACCATCTAC | CTGTCCGTTCAGGCTCTTCTT | Qiu et al., 2016 |
| *Per2* | TCCGAGTATATCGTGAAGAACG | CAGGATCTTCCCAGAAACCA | Kwapis et al., 2018 |
| *Adamts13* | AAACACTGCCTGATGTCCCG | GCACGGCAACCATAAGCAAA | custom-design |
